# Tracing the illicit Peruvian delicacy: proof-of-concept multiplex-PCR for the simultaneous detection of seven commercial *Macrobrachium* prawn species

**DOI:** 10.1101/2025.10.03.680321

**Authors:** Alan Marín, Ruben Alfaro, Cleila Carbajal, Lorenzo E. Reyes-Flores, Claudio Villegas-Llerena, Guillermo Mantilla, Karen Rodríguez-Bernales, Luis E. Santos-Rojas, Angel Yon-Utrilla, Eliana Zelada-Mázmela

## Abstract

The genus *Macrobrachium* (Decapoda, Palaemonidae) comprises nearly 300 prawn species distributed worldwide. At least nine of these species naturally inhabit the Pacific slope river systems of Peru, where most sustain significant artisanal fisheries and are highly prized as a luxury delicacy in the nation’s traditional cuisine. Here, we developed and validated a multiplex PCR assay with species-specific primers to identify seven *Macrobrachium* species of economic importance to Peru. Through robust validation, our assays achieved high specificity and efficiency, enabling fast and simultaneous identification of the seven target species. The utility of our novel multiplex PCR assay was evaluated as a proof-of-concept tool for conducting the first molecular monitoring of species diversity and illegal trade in Peruvian prawns across different commercial sectors. Among the 51 analyzed commercial prawn samples, a total of four species were detected, including *M. caementarius* (82.4%), *M. rosenbergii* (11.8%), *M. americanum* (3.9%), and *M. gallus* (1.9%). Notably, our molecular monitoring in restaurants revealed the first evidence of illegal trade in native wild-caught prawns in La Libertad region occurring during the reproductive closed season, when only the market of the exotic farmed prawn *M. rosenbergii* is permitted. These findings underscore the urgent need for enhanced monitoring programs in prawn fisheries in northern Peru, as current efforts are mainly focused on more productive central and southern regions. Our multiplex PCR assay provides a robust and efficient tool that can assist authorities and researchers in monitoring efforts, combating illegal trade, conducting diversity research, and supporting conservation.

## 1. Introduction

Peru is the second-largest producer of South American river prawns, following Brazil (FAO, 2024). In 2021, Peruvian aquaculture produced 25 tones of the exotic giant river prawn, *Macrobrachium rosenbergii*, valued at US$ 264,000 (FAO, 2024). Meanwhile, the harvest of the native freshwater prawn, *M. caementarius*, reached 298 tons, although no estimated value was available at the time of our analysis (FAO, 2024). The diversity of *Macrobrachium* species from the Peruvian Pacific slope includes at least nine nominal species: *M. americanum, M. caementarius, M. digueti, M. gallus, M. hancocki, M. inca, M. panamense, M. tenellum*, and *M. transandicum* (Luque, 2007; WoRMS, 2025; Zelada-Mázmela et al., 2025). Despite the high diversity of native Peruvian river prawns, they are commonly landed and marketed simply as “*camarón*” or “*camarón de río*” (Spanish for “prawn” and “river prawn”, respectively). The only prawn species whose production is registered with a specific market name is the exotic *M. rosenbergii*, which was introduced from Israel in 1980 (Miglio et al., 2021) and is marketed in Peru as “*camarón gigante de Malasia*” (Spanish for “Malaysian giant river prawn”) (PRODUCE, 2024).

Among the nine native *Macrobrachium* species found along the Peruvian Pacific slope, *M. caementarius* is the most harvested and widely traded across both coastal and Andean regions of Peru (Wasiw and Yépez, 2015). In contrast, the other species are primarily consumed locally or sold in nearby fishing grounds of central and northern coastal cities (Zelada-Mázmela et al., 2025). Two species, *M. hancocki* and *M. tenellum*, are rarely found, while a recent molecular phylogenetic study suggested the conspecific relationship between *M. digueti* and *M. transandicum* (Zelada-Mázmela et al., 2025). Consequently, only six native Peruvian prawn species (*M. americanum, M. caementarius, M. digueti, M. gallus, M. inca*, and *M. panamense*) are commonly harvested for domestic consumption.

Peruvian river prawns are considered a luxury delicacy, found in specialized local and high-end seafood restaurants known as “*cevicherías*” and “*picanterías*”. Their prices typically range from US$ 7 to 50 per kilogram, depending on factors such as the prawn’s size, the supply chain stage, the time of the year, and the location of sale. The high commercial value of Peruvian prawns, coupled with the so-called “Peruvian gastronomic boom” and a significant influx of tourists in cities during the summer holidays (from December to March) results in increased demand. This heightened demand and elevated prices during the Peruvian summer lead some fishermen and stakeholders to engage in illegal extraction and trade activities during the closed fishing season, which is enforced annually from January to March (R.M. N° 312-2006- PRODUCE; El Peruano, 2006). Evidence of this illegal trafficking is seen each year when Peruvian authorities confiscate large quantities of prawns during the closed season from different points of the supply chain, including fishing grounds, markets, and restaurants in the highest productive central and southern regions (Canal N, 2024). Moreover, to meet the high demand during the summer season, some restaurants may illegally mislabel wild prawns, which are extracted during the closed season, as the exotic farmed *M. rosenbergii*, for which no closed season applies (R.M. Nº 424-2014-PRODUCE; El Peruano, 2014).

Previous studies have reported clear signs of overexploitation in Peruvian prawn populations (Wasiw and Yépez, 2017; Pinazo et al., 2021). Peruvian authorities are attempting to recover these overfished populations through restocking programs using wild larvae sourced from various locations (ANDINA, 2024). Nevertheless, these programs are conducting without prior population genetic research, compromising the genetic integrity and long-term sustainability of these populations (Grant et al., 2017). To combat prawn overfishing and strengthen sustainability, the Peruvian prawn fishery is managed through regulations that include a minimum commercial size limit of 7 cm (R.M. Nº 209-2001-PRODUCE; El Peruano, 2001), a closed season during the breeding period of native wild *Macrobrachium* species (R.M. N° 312-2006-PRODUCE; El Peruano, 2006), and a policy that mandates fishermen to obtain official fishing permits. These permits allow them to harvest prawns using permitted artisanal fishing gear, but only from locations more than 5 km off the river mouth (R.M. Nº 083-2007- PRODUCE; El Peruano, 2007).

Notwithstanding the existing protective measures, several management flaws need to be addressed. For example, current conservation efforts including periodic stock assessments and monitoring, aquaculture, and biological research primarily focus on a single species: *M. caementarius* (Bazán et al., 2009; Reyes, 2014; Wasiw and Yépez, 2015; Zacarías-Ríos and Yépez-Pinillos 2015; Flores-Gómez, 2021, Reyes, 2019). On the other hand, the other native species receive significantly less attention, and only few studies have been published about them, focusing on their taxonomy (Méndez, 1981), aquaculture (Dávila et al., 2013), and molecular systematics (Zelada-Mázmela et al., 2025). Excepting *M. caementarius*, landing records for the other native prawn species are only recorded at the genus level (PRODUCE, 2024), affecting stock estimates necessary for effective species-specific management programs (Marín et al., 2022). Accurate identification of commercially important prawn species is a critical step required for proper management and decision-making process, particularly when the commercial catch comprises several closely related sympatric species that exhibit high morphological similarity, as is the case with Peruvian *Macrobrachium* species.

The strong interspecific conservatism and intraspecific plasticity observed in several *Macrobrachium* species hinder their species discrimination based on conventional morphological methods (Pileggi and Mantelato, 2010; Rossi and Mantelato, 2013; Zelada-Mázmela et al., 2025). Moreover, the diagnostic features for identifying *Macrobrachium* species described in most morphological keys primarily focus on adult male individuals (Méndez, 1981; Valencia and Campos, 2007), which typically possess a well-developed and distinctive second pair of chelated pereiopods (Thiel and Watling, 2015). Consequently, identifying females and juvenile *Macrobrachium* prawns can be challenging, even for trained experts. Furthermore, species-level identification is more problematic or even entirely prevented in highly processed samples and cooked presentations, as many key features are removed or significantly altered during these processes.

In this context, DNA-based methods are highly effective for accurate species identification, even when dealing with highly processed seafood samples (Marín et al., 2018, 2025; Hellberg, 2024; Marín, 2025). Among these methods, DNA sequencing stands out as one of the most accurate techniques for identifying prawn species (Pileggi and Mantelato, 2010; Rossi and Mantelato, 2013; Zelada-Mázmela et al., 2025). However, there is a pressing need to develop and validate faster and cheaper DNA-based assays to complement conventional monitoring methods, support aquaculture activities, and advance biological research. Species-specific primers (SSPs) combined with endpoint multiplex Polymerase Chain Reaction (PCR) is both time-efficient and cost-effective approach. This method enables accurate and simultaneous species discrimination, even among closely related ones (Damasceno et al., 2016; Wilwet et al., 2021; Marín et al., 2025).

Despite the high diversity of Peruvian *Macrobrachium* species that hold both ecological and commercial significance, the taxonomic differentiation of these species presents challenges. To our knowledge, no study has yet developed species-specific DNA-based methods, such as SSP assays, for identifying *Macrobrachium* species from the Peruvian Pacific slope. Therefore, this study aims to develop and apply a multiplex PCR-SSP assay targeting the simultaneous identification of seven *Macrobrachium* species of commercial value within the Peruvian seafood sector. Our molecular identification protocol was standardized and validated using DNA from vouchered specimens and applied as a proof-of-concept tool for authenticating and monitoring prawn products collected in three important sectors: 1) the seafood sector, to monitor the illegal trade of cooked samples collected during the closed fishing season; 2) the aquaculture sector, to authenticate farmed prawn species; and 3) the pet shop aquarium sector.

## 2. Materials and methods

### 2.1 Collection of fresh prawn samples and DNA extraction

The genetic material of six native prawn species (*M. americanum, M. caementarius, M. digueti, M. gallus, M. inca*, and *M. panamense*) used in our molecular validation assays were sourced from vouchered specimens collected along the Peruvian Pacific slope. They were identified both morphologically and molecularly using a DNA barcoding approach, as detailed in a recent study (Zelada-Mázmela et al., 2025). For more information regarding the collection procedures, sampling sites, DNA extraction, and species identification methods, please refer to the aforementioned reference.

### 2.2 Collection of commercial prawn samples and DNA extraction

Commercial samples used to assess the effectiveness of our SSP identification protocols included live, fresh, and cooked presentations. These samples were collected between April 2024 and May 2025 using non-probabilistic purposive sampling. Aquatic and seafood businesses were chosen for featuring prawn-related advertisements on their social media platforms. A live prawn specimen was acquired from a pet shop aquarium in La Libertad region. Exotic *M. rosenbergii* specimens were sourced from an aquaculture facility in the Ancash region. Fresh and cooked samples were obtained throughout the year during both open and closed fishing seasons, from a gastronomic festival and restaurants in Tumbes, La Libertad, Ancash, Lima, and Arequipa regions (see Table 1 and Fig. 1).

**Table 1.**
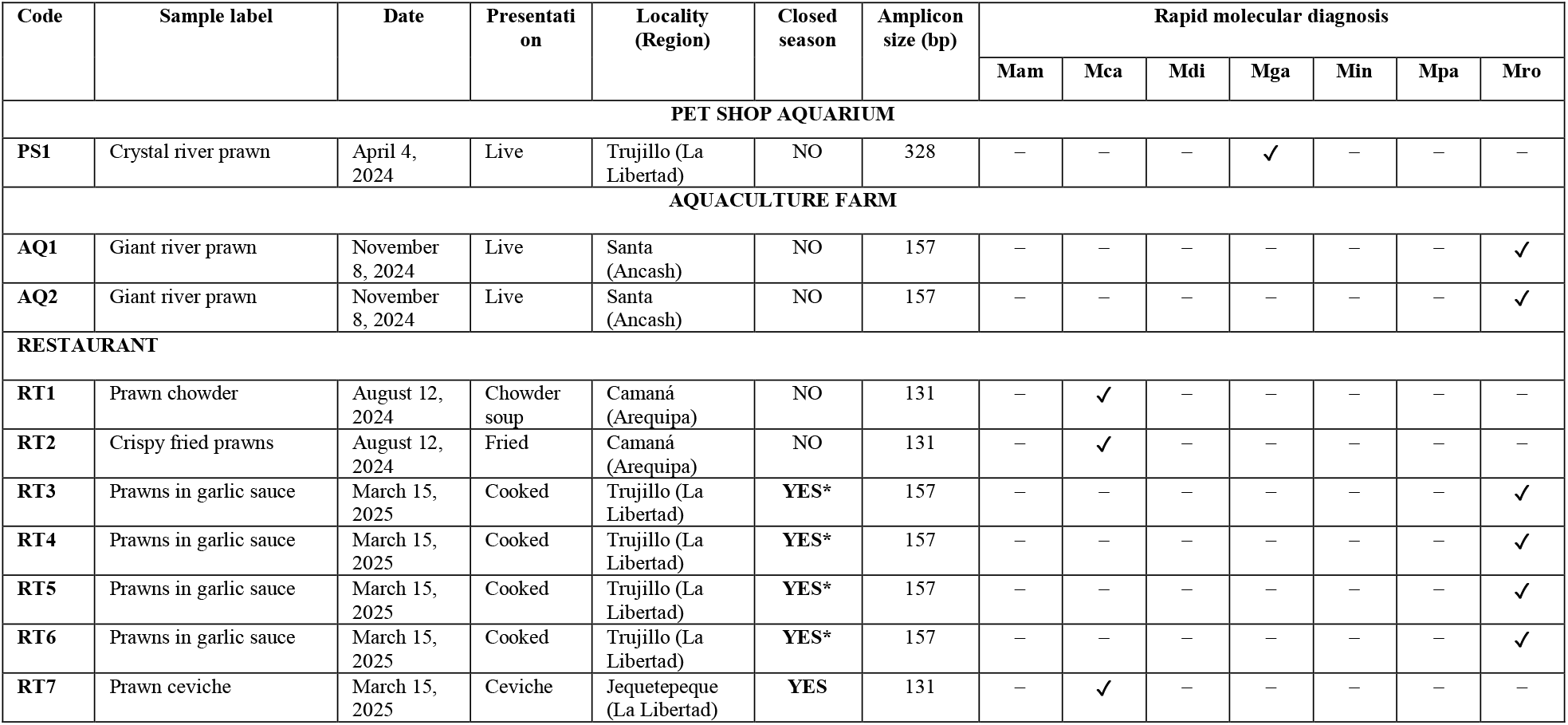
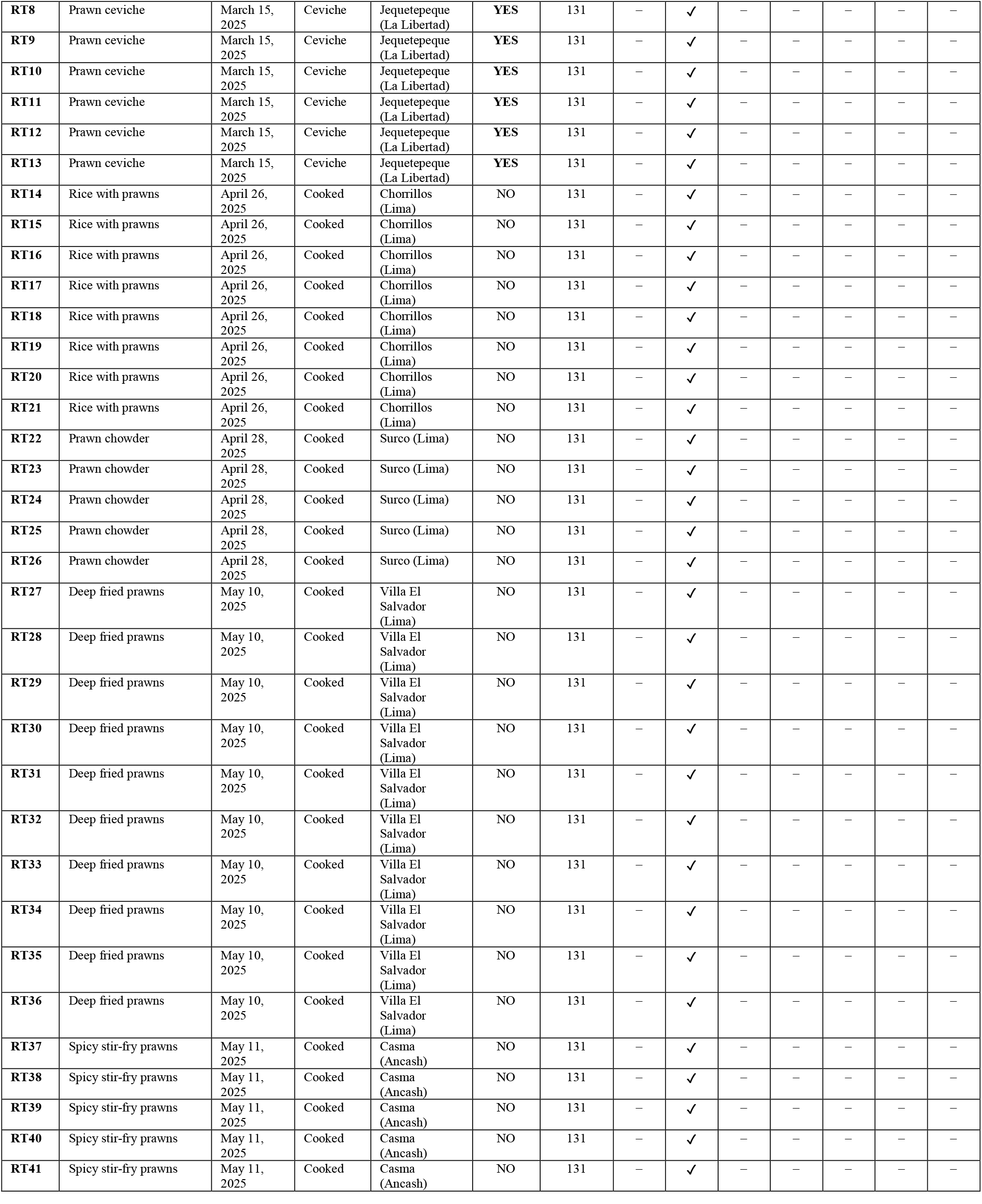
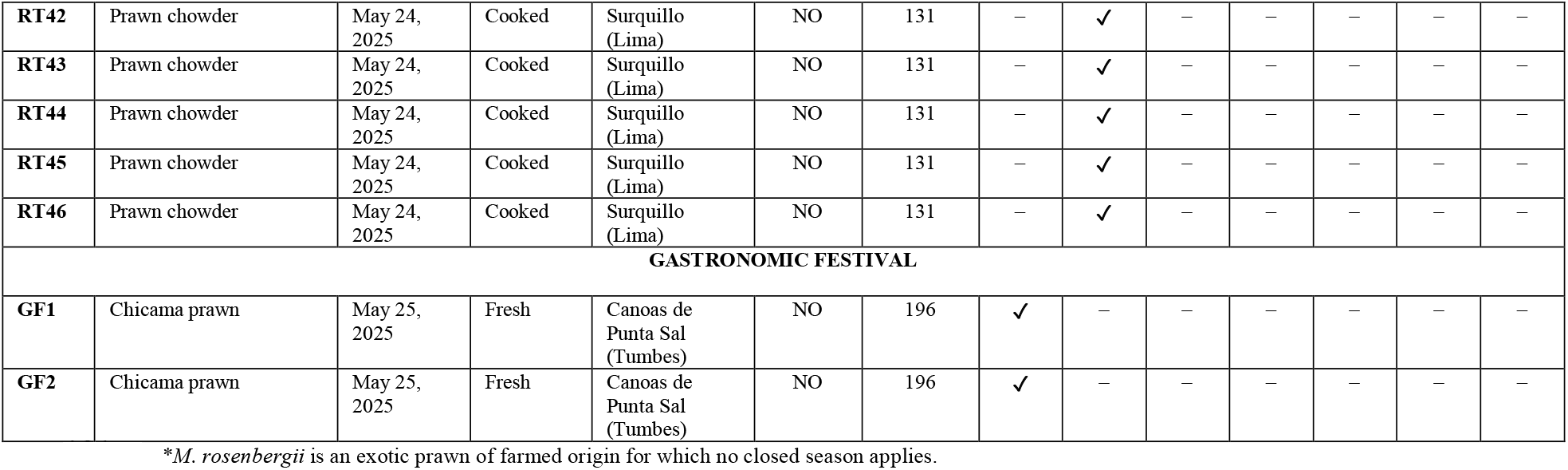
River prawn samples of commercial origin used in the authentication multiplex-PCR assay with the species-specific primers developed in this study. Mam: *Macrobrachium americanum*, Mca: *M. caementarius*, Mdi: *M. digueti*, Mga: *M. gallus*, Min: *M. inca*, Mpa: *M. panamense*, Mro: *M. rosenbergii*.

**Fig. 1.**
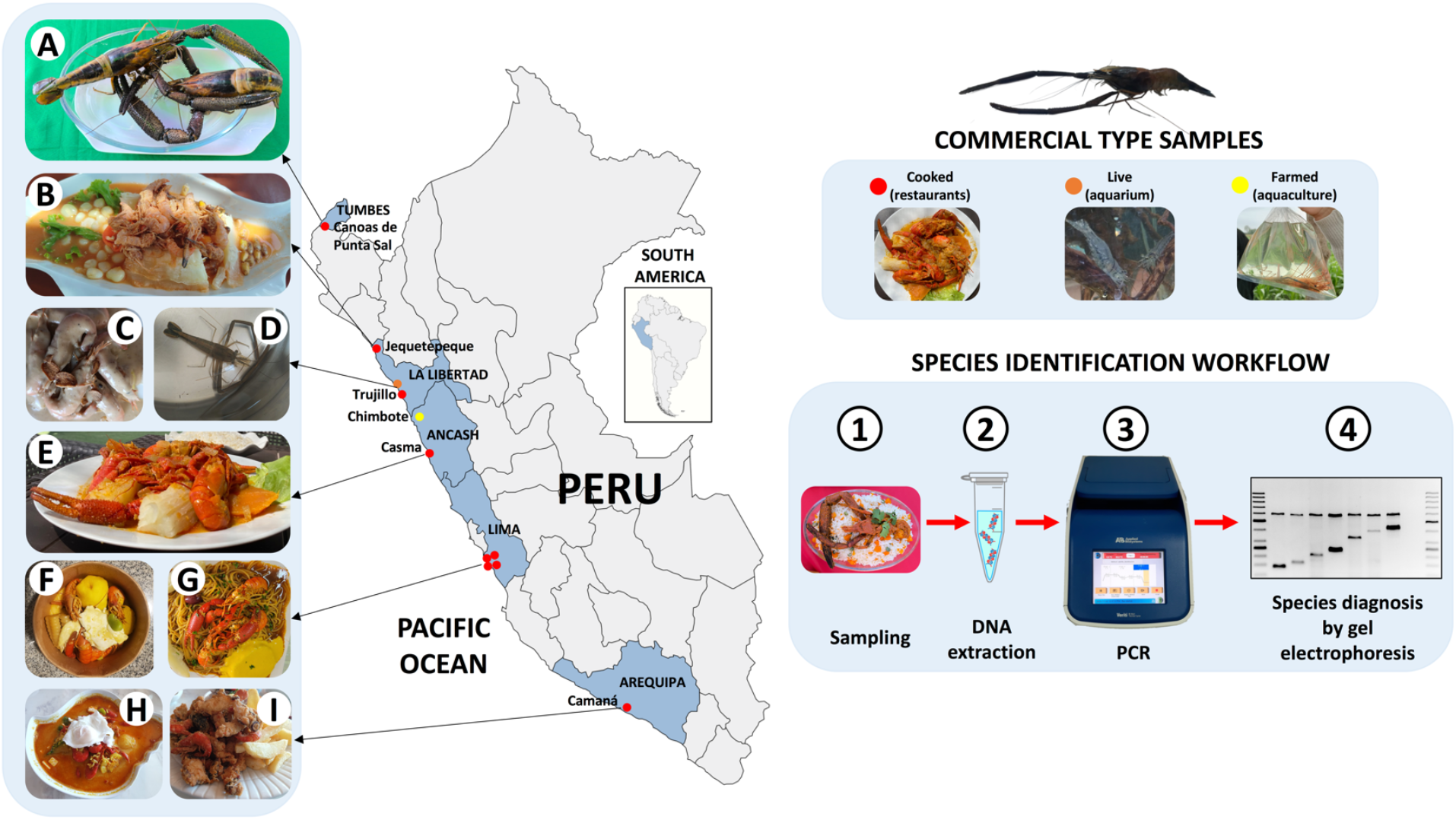
Prawn sampling map. Peruvian sampling locations of the commercial river prawns analyzed in this study. Panel A fresh sample collected in Tumbes region; Panel B cooked sample collected in Jequetepeque city; Panel C cooked sample collected in Trujillo city; Panel D live sample from a pet shop aquarium located in Trujillo city; Panel E cooked sample collected in Casma city; Panels F and G cooked samples collected in Lima region; and Panels H and I cooked samples collected in Arequipa region.

From each sampled specimen, one chela was collected, rinsed with distilled water, and stored individually in a Falcon tube containing 96% ethanol. In cooked prawn samples, the chela was chosen from whole organisms, and any detached prawn parts were excluded. This practice not only prevented cross-contamination but also ensured that each individual was uniquely identified, avoiding redundant testing. The live prawn specimen was cold- anesthetized and preserved in 96% ethanol prior to dissection. All collected samples were stored at -20°C until DNA analysis. Total genomic DNA (gDNA) was isolated using the standard phenol- chloroform protocol (Sambrook and Russell, 2001). DNA quantification was performed using an Epoch spectrophotometer (BioTek Instruments, Winooski, VT, USA). The total gDNA was diluted to a final working concentration of 20 ng/µl and stored at -20°C for further PCR analyses.

### 2.3 Primer design

#### 2.3.1 Design of forward common and reverse species-specific primers

The mitochondrial *cytochrome oxidase subunit I* (COI) gene was screened for species-specific single nucleotide polymorphisms (SNPs) in the seven target prawn species: *M. americanum, M. caementarius, M. digueti, M. gallus, M. inca, M. panamense*, and *M. rosenbergii*. The COI gene was selected based on recent research on the phylogenetics of Peruvian *Macrobrachium* species (Zelada-Mázmela et al., 2025), which indicated that the COI gene sequences exhibit the greatest interspecific genetic distances compared to 16S rRNA and 28S rDNA genes. For the design of SSPs, available DNA sequences from the 5’ end of the COI gene for the seven target *Macrobrachium* species were retrieved from the BOLD and GenBank databases. The data included 360 DNA sequences for *M. rosenbergii*, 86 sequences for *M. americanum*, 45 sequences for *M. inca*, 41 sequences for *M. caementarius*, 16 sequences for *M. digueti*/*M. transandicum*, 7 sequences for *M. gallus*, and 4 sequences for *M. panamense*. All DNA sequences were then multialigned using MEGA 7 software (Kumar et al., 2016).

A common forward primer, designated MacrCOIU-F, was designed based on a conserved region detected at the 5’ end of the COI gene across all analyzed *Macrobrachium* species (see Fig. 2). The design of the SSPs followed the methodologies outlined by Wang et al. (2018) and Marín et al. (2025). For each target *Macrobrachium* species, three potential reverse SSPs were manually designed based on species-specific SNPs located at the 3’-terminus (first nucleotide from the 3’ end) of each SSP. Having a mismatch at the 3’-terminus of the SSP compared to a homologous region in non-target species can disrupt polymerase activity at their hybridization, thereby ensuring PCR amplification is specific to the target species’ locus (Stadhouders et al., 2010). In some instances, it was necessary to intentionally introduce of a single mismatch at the second, third, or fourth nucleotide position from the primer’s 3’ end to guarantee absolute allelic specificity for the target species (Wang et al., 2018).

**Fig. 2.**
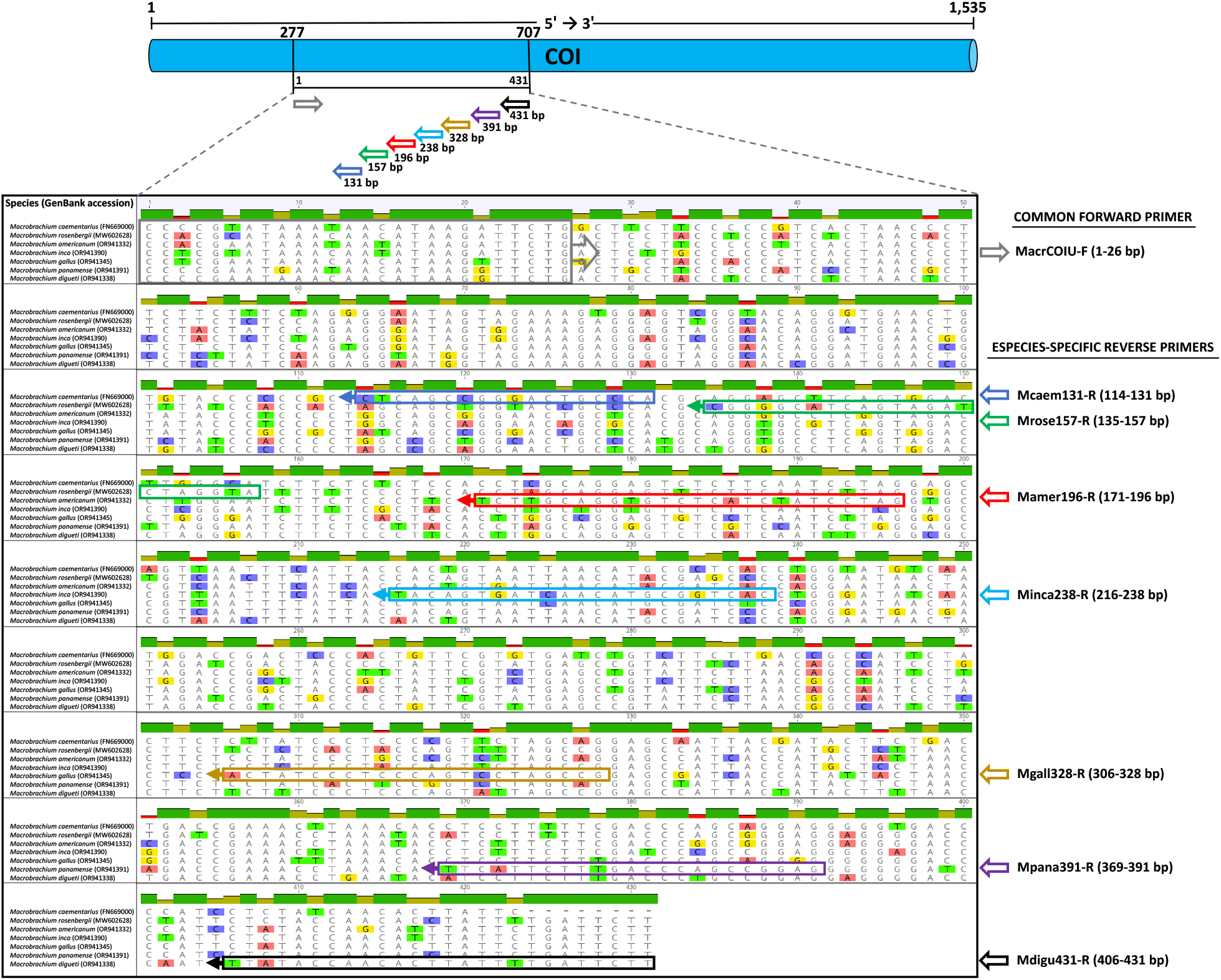
Schematic representation of species-specific primer design for the seven *Macrobrachium* species targeted in this study.

Additionally, a deliberately “GG tail” was added at the 5’ terminus of some SSPs to meet the minimum percentage of recommended GC content (40%). This enhancement ensures more stable primer/template binding during PCR amplification (Lu et al., 2018; Marín et al., 2025). The combination of the forward common primer MacrCOIU-F with each reverse SSP was designed to yield amplicon lengths of different sizes ranging from 100 to 500 bp for each target species. This small amplicon size range facilitates effective PCR amplification in cooked samples, where gDNA integrity and quality is compromised due to degradation from physical and chemical processes (Marín, 2025; Marín et al., 2025). To maintain a high astringency in PCR reactions, all primers were designed to work at high annealing temperatures of 60°C or above. A summary of all SSPs designed in this study are presented in Table 2.

**Table 2.**
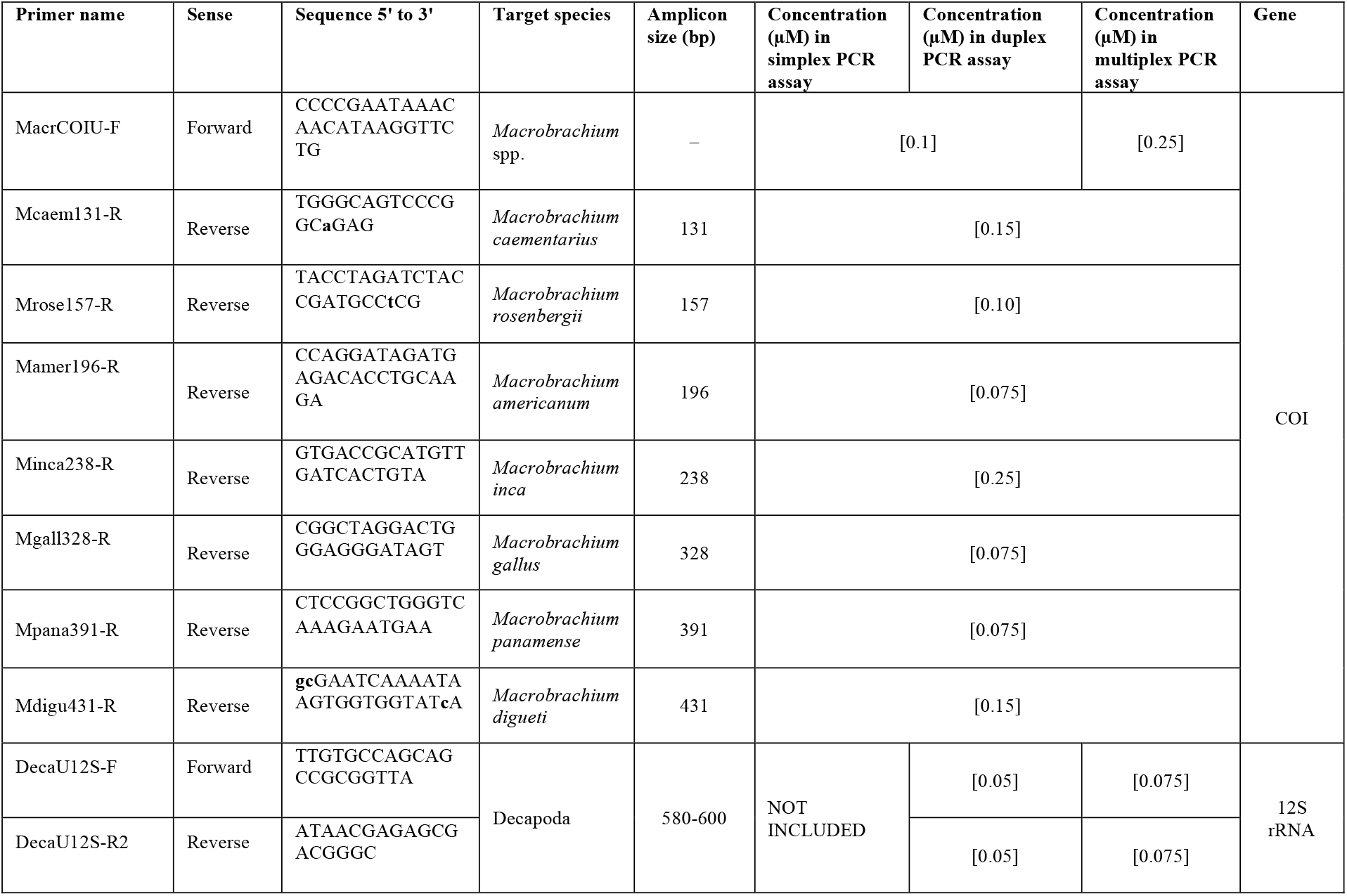
List of species-specific and common primers developed in this study for the identification of river prawn species from the genus *Macrobrachium*. Deliberate nucleotide mismatches introduced in primer sequences are written in lowercase and highlighted in bold.s.

#### 2.3.2 Design of internal endogenous control primer set

An internal endogenous control primer set, designated DecaU12S-F/R (refer to Table 2), was developed to target a partial fragment (580-600 bp) within the mitochondrial 12S rRNA gene of the analyzed *Macrobrachium* species. This fragment size range was chosen because it exceeds the length of the amplicons generated by the SSPs, facilitating straightforward differentiation during analysis. This endogenous control primer set was designed based on two highly conserved regions within the 12S rRNA gene sequences of 14 Decapoda species that were obtained from their complete mitogenomes retrieved from the GenBank database. These species were: *Cherax quadricarinatus* (GenBank access MN648457), *Lithodes nintokuae* (GenBank access NC_024202), *Macrobrachium rosenbergii* (GenBank access MZ781226), *Metacarcinus magister* (GenBank access OR400215), *Metapenaeus affinis* (GenBank access ON599338), *Neocaridina denticulata* (GenBank access MW238411), *Panulirus longipes* (GenBank access NC_052749), *Penaeus (Farfantepenaeus) californiensis* (GenBank access NC_012738), *Penaeus (Litopenaeus) stylirostris* (GenBank access MN879254), *Penaeus (Litopenaeus) vannamei* (GenBank access KT596762), *Penaeus (Penaeus) monodon* (GenBank access NC_002184), *Pleoticus muelleri* (GenBank access NC_03996), *Solenocera melantho* (GenBank access NC_06904), and *Trachysalambria curvirostris* (GenBank access MK681887).

### 2.4 In-silico analysis

The potential formation of secondary structures such as hairpins, homo- and hetero-dimers, were evaluated using two independent online tools: the IDT OligoAnalyzer tool (available at <http://www.idtdna.com>) and The Sequence Manipulation Suite (Stothard, 200) (available at <http://www.bioinformatics.org/sms2/index.html>). Additionally, an *in-silico* analysis was performed to verify the specificity of each SSP using the Primer-BLAST tool (Ye et al., 2012). For this analysis, the “nt” nucleotide database was selected, and organisms were limited to those within the Infraorder Caridea (taxid: 6694). Any matches to Caridean species not found in rivers of the Peruvian Pacific slope were disregarded. The newly designed oligonucleotides were synthesized by Integrated DNA Technologies (IDT, Coralville, Iowa, USA) and Invitrogen (Carlsbad, CA, USA).

### 2.5 Simplex PCR-SSP assay standardization

The specificity of each SSP set was initially assessed using gradient PCR analyses on two individuals from each target and non-target *Macrobrachium* species. These analyses evaluated annealing temperatures ranging from 60 to 64°C at 1°C intervals and primer concentrations between 50 and 500 nM, increasing in increments of 25 nM. Subsequently, the optimal standardized simplex PCR-SSP conditions were applied to amplify the gDNA from all collected individuals of each target species. All PCR analyses conducted in this study (simplex, duplex, and multiplex) were performed in a Veriti 96 Well thermal cycler (Applied Biosystems, Foster City, CA, USA) with the Maximo *Taq* DNA Polymerase 2x-preMix (GeneOn GmbH, Nürnberg, Germany). The standardized simplex PCR-SSP was performed under the following thermal cycling conditions: an initial denaturation at 94°C for 5 min, followed by 35 cycles of 95°C for 15 s, 60°C for 15 s, and 72°C for 15 s, concluding with a final extension step at 72°C for 3 min. The PCR cocktails consisted of a final volume of 10 µl, containing 5 µl of Maximo *Taq* DNA Polymerase 2x-preMix (GeneOn GmbH, Nurnberg, Germany), 0.1 µl of the common forward primer MacrCOIU-F (10 µM), the standardized concentration for each SSP (as indicated in Table 2), 0.5 µl of template DNA (20 ng/µl), and filled up with ultrapure H_2_O to reach the final volume of 10 µl. Three µl of each PCR reactions were electrophoresed along with a 50 bp molecular ladder (SIGMA) on a 2% agarose gel (EMD Millipore, Billerica, MA, USA), stained with GelRed Nucleic Acid Gel Stain (BIOTREND Chemikalien GmbH, Köln, Germany), and visualized in an UVIDOC HD6 Gel Documentation System (UVITEC Cambridge, UK).

### 2.6 Duplex PCR-SSP assay standardization

For the interspecific validation of the SSP against non-target species, a duplex PCR-SSP assay was standardized. This assay included the common forward primer MacrCOIU-F, a reverse SSP, and the novel internal endogenous control primer set DecaU12S-F/R (Table 2). In the absence of target DNA, non-target prawn species were expected to yield only the common internal endogenous control locus (12S rRNA). This would result in the visualization of a single band of about 580-600 bp on the agarose gel. In contrast, target prawn species were expected to produce two bands: one corresponding to a diagnostic species-specific locus (COI), generated by the combination of the forward MacrCOIU-F primer with each reverse SSP, and a second amplicon of higher molecular weight corresponding to the common internal endogenous control (12S rRNA). The final duplex PCR amplification protocol and master mix preparation were the same as used in the simplex PCR assay described before, with the addition of the internal endogenous control primer set DecaU12S-F/R at a final concentration of 0.05 µM for each primer in a total reaction volume of 10 µl. A 50 bp molecular ladder (SIGMA) was electrophoresed along the PCR products for size determination on a 2% agarose gel electrophoresis.

### 2.7 Multiplex PCR-SSPs assay standardization

The final multiplex PCR-SSPs assay included the common forward primer MacrCOIU-F, along with the seven reverse SSPs and the internal endogenous control primer set DecaU12S- F/R. The total volume for the multiplex reaction was 10 µl. The PCR amplification cycling protocol and primer concentrations were the same used in the duplex assays, with two minor adjustments: the concentration of the common forward primer MacrCOIU-F was increased to 0.25 µM, and the concentration of each internal endogenous control primers DecaU12S-F/R was set at 0.75 µM (refer to Table 2). All PCR products were visualized using a 2% agarose gel electrophoresis, with 3 µl of each amplicon loaded onto the gel. A 50 bp molecular ladder (SIGMA) was run alongside the PCR products for size determination. Successful PCR reactions were expected to yield two distinct bands: one representing the species-specific locus (COI), with size varying by prawn species, and another for the internal endogenous control locus (12S rRNA), approximately 580-600 bp in length.

### 2.8 Authentication of commercial prawn samples through multiplex PCR-SSPs

To evaluate the performance of our standardized multiplex PCR assay for authenticating prawn samples of commercial origin, we used the gDNA isolated from samples collected from an aquarium shop, an aquaculture facility, a gastronomic festival, and restaurants located along the Peruvian coast (see Table 1 and Fig. 1). The PCR amplification conditions and cocktail mix preparation were consisted with the procedures described in the previous section 2.7. These results enabled us to monitor not only the diversity of economically valuable prawn species in Peru but also instances of illegal trade involving prawn samples collected during the closed season. During this period, only the trade of farmed *M. rosenbergii* is allowed by Peruvian regulations of prawns’ fishery.

## 3. Results and discussion

We successfully developed and applied a novel multiplex PCR-SSP assay for the simultaneous identification of seven commercially important species of Peruvian river prawns from the genus *Macrobrachium*. To the best of our knowledge, this assay is the first of its kind focused solely on identifying *Macrobrachium* species. Our molecular assay demonstrated high accuracy and effectiveness in authenticating both fresh and processed prawn samples of commercial origin. This capability allowed us to monitor the trade of cooked prawns collected from restaurants during the closed season.

Prior studies on the simultaneous identification of *Macrobrachium* species using DNA-based technologies include methods such as PCR-RAPD (Meruane et al., 1996), PCR-RFLP (Aoki et al., 2013), DNA barcoding (Liu et al., 2007; Rossi and Mantelato, 2013; Siriwut et al., 2021), and DArT markers (Makombu et al., 2019). While these molecular methods have proven effective for identifying *Macrobrachium* species, they come with certain disadvantages that should be considered. For example, the reproducibility of PCR-RAPD can be affected by slight changes in the quality and quantity of target DNA (Yamashita et al., 2024). PCR-RFLP assays can be time-consuming due to the incubation step required for proper enzymatic digestion, and incomplete digestion may lead to misleading diagnostic results (Marín, 2025). DNA barcoding and DArT markers tend to be expensive, time consuming, and require specialized equipment, dedicated software, and trained personal (Zhou et al., 2015; Benamozig et al., 2021; Marín, 2025).

On the other hand, SSP identification assays are both cost- and time-effective, requiring only basic equipment and expertise. The main shortcomings of this approach occur during the development of the SSPs. Designing these primers requires bioinformatics knowledge and access to a large number of DNA sequences from both target and non-target species. A key aspect of designing successful SSPs is detecting species-specific SNPs within candidate genes, which can be particularly challenging among closely related and congeneric species (Marín, 2025; Marín et al., 2025).

### 3.1 Species-specific primers design and performance

For SSP design, we utilized a total of 559 COI gene sequences from the seven target species. The analyzed sequences exhibited some species-specific SNP positions in which we anchored the 3’-terminus (first nucleotide from the 3’ end) of each candidate SSP. These specific positions were crucial for destabilizing the hybridization of the SSPs to homologous regions in non-target prawn species, thereby ensuring that the amplification of the COI locus was specific to each target species and preventing cross-species reactions. In addition, the use of modern PCR kits and master mixes containing a pre-formulated buffer with ammonium sulfate, such as the commercial kit used herein, greatly increases the specificity of the SSPs (Marín et al., 2025). The ammonium ion destabilizes the weak hydrogen bonds present in mismatched primer-template bases, thus optimizing specificity by preventing primers from binding to non-specific sites (Thermo Fisher Scientific, n.d.). The locations of the hybridization sites for each SSP within the mitochondrial COI gene are depicted in Fig. 2, while the nucleotide sequences and the expected amplicon sizes for each SSP are provided in Table 2.

The successful performance of the SSPs was initially predicted through robust *in-silico* tests. The results of the *in-silico* specificity validation analysis, conducted using the Primer Blast software, demonstrated that all SSPs exhibited the highest affinity solely for their respective target species. This performance was subsequently confirmed through three *in-vitro* validation assays including simplex (Panels 1A to 7A in Fig. S1), duplex (Panels 1B to 7 B in Fig. S1), and multiplex PCR (Fig. 3), using the DNA of fresh and cooked specimens.

**Fig. 3.**
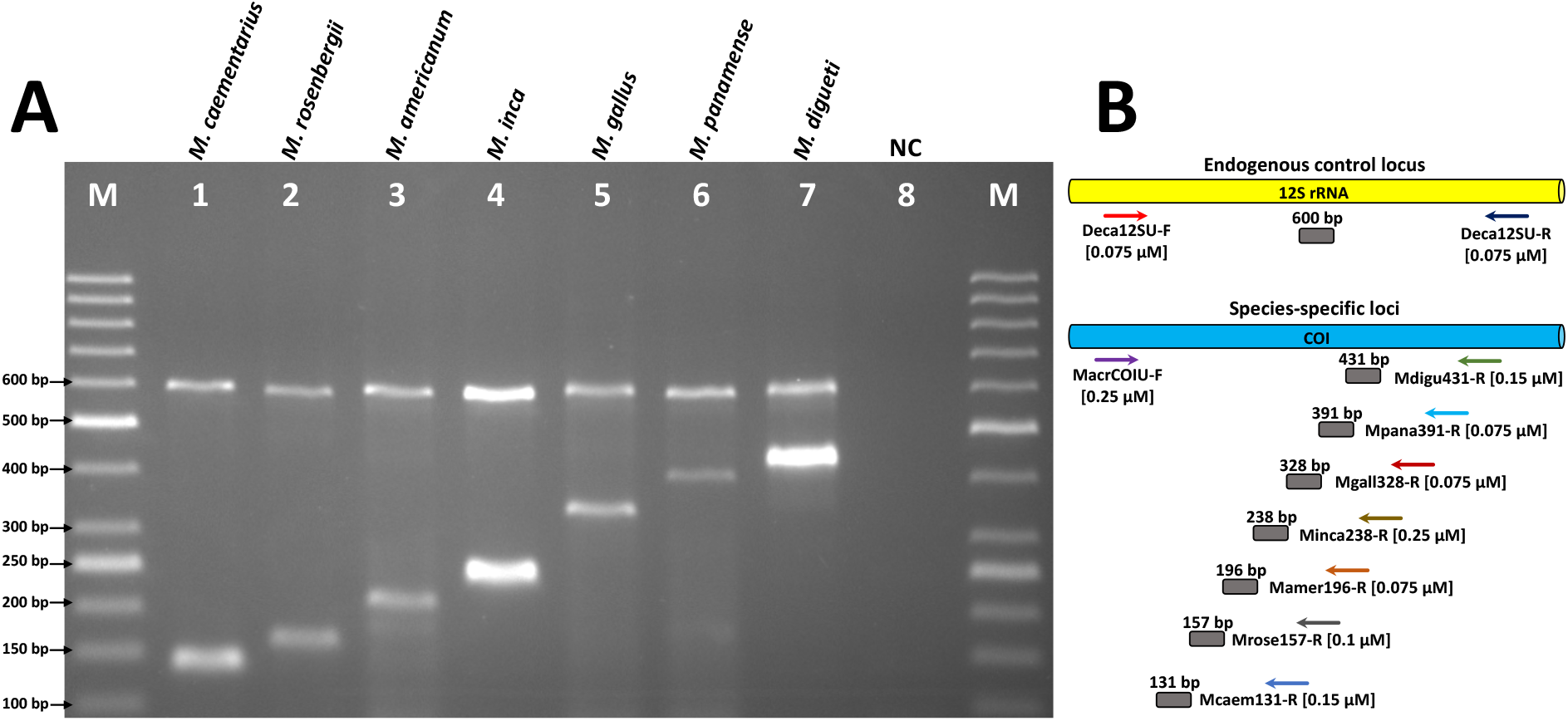
Multiplex validation results. Panel A shows the agarose gel with the multiplex PCR results using the species-specific primers targeting seven *Macrobrachium* species. Panel B shows and schematic representation of the positions for each species-specific primer and the endogenous control primer set, their expected amplicon sizes, and primer concentrations. M: molecular ladder (50 bp) loaded into wells at both ends of the agarose gels. NC: negative control.

To enhance the specificity of SSPs that were designed over regions containing few species-specific SNPs, we intentionally introduced mismatches at specific nucleotide positions from these primer sequences. For the SSP Mcaem131-R, which targets *M. caementarius*, a T→A mismatch was added at the fourth nucleotide position. Similarly, a C→T mismatch was introduced at the third nucleotide position of SSP Mrose157-R, targeting *M. rosenbergii*, and a A→C mismatch was added at the second nucleotide position of SSP Mdigu431-R, which targets *M. digueti*. These three SSPs with mismatches showed greater specificity compared to the candidate SSPs without introduced mismatches that were designed over the same regions targeting these three species. Importantly, the deliberately introduced mismatches did not compromise PCR amplification of DNA from their respective target species, as evidenced by the results of the *in-vitro* validation assays (Fig. S1 and Fig. 3). This primer design strategy, involving deliberate mismatches at the 3’-terminus, has been shown effective in previous studies (Wang et al., 2018 and references therein).

### 3.2 Simplex PCR-SSP intraspecific validation

Our simplex PCR assays utilized the common forward, MacrCOIU-F, along with a species-specific reverse primer. Each simplex assay successfully amplified the DNA from all tested specimens. These results confirm the intraspecific affinity of each SSP, as shown in the agarose gel electrophoreses from Panels 1A to 7A of Fig. S1. A specific PCR product of the expected size was obtained for each target species, as follows: *M. caementarius* with 131 bp (Panel 1A in Fig. S1), *M. rosenbergii* with 157 bp (Panel 2A in Fig. S1), *M. americanum* with 196 bp (Panel 3A in Fig. S1), *M. inca* with 238 bp (Panel 4A in Fig. S1), *M. gallus* with 328 bp (Panel 5A in Fig. S1), *M. panamense* with 391 bp (Panel 6A in Fig. S1), and *M. digueti* with 431 bp (Panel 7A in Fig. S1).

### 3.3 Duplex PCR-SSP interspecific validation

The second validation step involved a duplex PCR, which utilized the common forward primer, MacrCOIU-F, a reverse SSP, and the endogenous control primer set namely DecaU12S-F/R. This validation aimed to assess the specificity of each SSP against the DNA from non-target *Macrobrachium* species. During this validation, no cross-species reactions were observed. DNA from the non-target species produced only the band for the endogenous control (12S rRNA gene) at approximately 580-600 bp (see Panels 1B to 7B in Fig. S1). These results demonstrate the high specificity of the SSPs and the successful enzymatic reaction during the PCR.

### 3.4 Multiplex PCR-SSP validation

The final *in-vitro* validation assay involved an octuplex PCR that enabled the simultaneous amplification of the internal endogenous 12S rRNA control locus along with the seven SSP loci from the COI gene (Fig. 3). Each SSP locus generated a unique fragment size, facilitating easy species discrimination by comparing PCR product sizes against a 50 bp DNA ladder on an agarose gel. Our seven SSPs demonstrated high effectiveness under the high-stringency conditions of the multiplex PCR, yielding unambiguous specific amplicons at an annealing temperature of 60°C. Moreover, the short time allocated for each of the three PCR steps (denaturation, hybridization, and elongation was 15 s each) allowed for a total PCR cycling analysis of about 50 min. Combining this with a 30-min electrophoresis run, we achieved fast species identification results within 1 h and 20 min. This time estimate does not include the durations for DNA extraction and master mix preparation, which can vary depending on the protocol and sample size.

### 3.5 Proof-of-concept: Application of the multiplex PCR-SSP assay to monitor the trade of commercial prawn samples

The applicability and performance of our standardized multiplex PCR assay was further evaluated using commercial prawn samples of various types (live, fresh, and cooked) collected from a pet shop aquarium (n=1), an aquaculture facility (n=2), a gastronomic festival (n=2), and several restaurants (n=46) (Table 1 and Fig. 1). Our multiplex assay achieved 100% efficiency and accuracy in identifying these 51 commercial samples to the species-level. Overall, four species were detected, including *M. caementarius* (82.4%), *M. rosenbergii* (11.8%,), *M. americanum* (3.9%), and *M. gallus* (1.9%). The sample collected from the pet shop aquarium was accurately identified as *M. gallus* (see Panel D in Fig. 1, sample code PS1 in the agarose gel of Fig. 4), while those from the aquaculture facility were confirmed as *M. rosenbergii* (sample codes AQ1 and AQ2 in the agarose gel of Fig. 4). Two fresh prawn samples obtained from a gastronomic festival held in Canoas de Punta Sal district (Tumbes region) were identified as *M. americanum* (see Panel A in Fig. 1, sample codes GF1 and GF2 in the agarose gel of Fig. 4). Additionally, we collected 46 cooked prawn samples from eight restaurants in the following districts: Jequetepeque and Trujillo (La Libertad region), Casma (Ancash region), Chorrillos, Surco, Surquillo, and Villa el Salvador (Lima region), and Camaná (Arequipa region) (Table 1). Among the 46 cooked samples, 42 were identified as *M. caementarius* (91.3%) and four as *M. rosenbergii* (8.7%,). These results demonstrate that our multiplex assay can be effectively used to monitor the prawn species-composition and trade in restaurants. This is important because we demonstrated that our novel detection assay can be used to support monitoring efforts and combat the illicit prawn trade in restaurants.

**Fig. 4.**
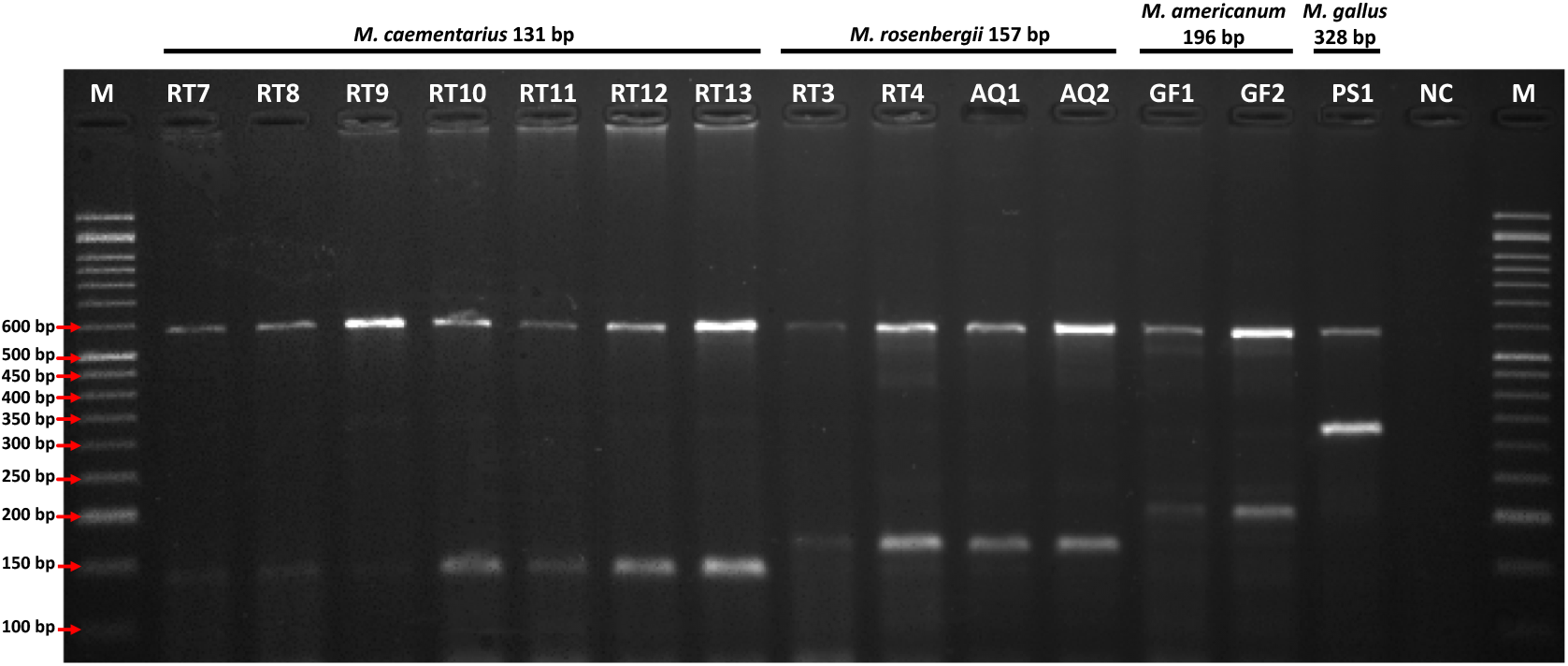
Multiplex-PCR authentication of prawn samples collected from restaurants, an aquaculture facility, a gastronomic festival, and a pet shop aquarium. M: molecular ladder (50 bp) loaded into wells at both ends of the agarose gels. NC: negative control. For the sample codes refer to Table 1.

During the ban on native river prawn fishery and trade, two seafood restaurants in northern Peruvian cities namely Jequetepeque and Trujillo (La Libertad region) were found offering river prawns on their menus. Seven prawn ceviche samples collected in the restaurant from Jequetepeque city were clearly identified by our multiplex-PCR assay as *M. caementarius*, as revealed by the visualization of amplicons measuring 131 bp (see Panel B of Fig. 1, sample codes RT7 to RT12 in Table 1 and the gel electrophoresis in Fig. 4). According to the restaurant staff, these samples were obtained from artisanal fishermen who caught the prawns from a local river, presumably the Jequetepeque river. This result represents the first evidence on illegal trade in wild-caught prawns during the closed season in this city. Despite the lack of recent scientific surveys, local ecological knowledge from the Jequetepeque river basin suggests a significant decline in prawn populations (Gálvez-Mora, 2022). These findings underscore the urgent need for prawn fishery monitoring programs and population assessments in the Jequetepeque river basin. To curb illegal trade during the closed season, improved traceability and monitoring systems are needed across the prawn supply chain in Jequetepeque city and its neighboring towns.

The other restaurant offering prawns during the closed season is located Trujillo city and featured a menu item labeled “prawn in garlic sauce”. After conducting the multiplex PCR analysis on these cooked samples, we observed PCR products measuring 157 bp, which confirmed that these samples were the giant river prawn, *M. rosenbergii* (refer to Panel C in Fig. 1, sample codes RT3 to RT6 in Table 1 and gel electrophoresis in Fig. 4). As mentioned earlier, this species is not subject to closed fishing regulations, verifying the legal commercial origin of these samples. Under Peruvian Law R.M. Nº 424-2014-PRODUCE (El Peruano, 2014), the exotic *M. rosenbergii* is the only prawn that can be marketed throughout the year, as it is produced by the aquaculture sector. Consuming farmed prawns presents a legal and reliable alternative that can help alleviate the high demand and reduce the illegal trade of wild-caught prawns during closed seasons.

As anticipated, most of the restaurant samples examined were identified as *M. caementarius*, which is the most traded prawn in the country. In our assay, we specifically designed the SSP Mcaem131-R to target *M. caementarius* and produce a small-sized amplicon of 131 bp. This approach was adopted to ensure successful PCR amplification, even in the presence of degraded DNA obtained from processed or cooked samples collected from restaurants. Research has shown that cooked seafood samples exposed to high temperatures (such as boiling and deep-frying) and low pH levels (as seen in marinated ceviche) can lead to the degradation of gDNA. This degradation negatively affects downstream applications and hampers PCR amplification (Marín et al., 2018). Our SSPs successfully authenticated all cooked samples, regardless of their presentation (including ceviche, deep-fried, stir-fried dishes, and chowder). These results demonstrated the high efficiency and accuracy of our method in identifying cooked prawn samples exhibiting some extent of DNA damage.

*M. caementarius* is the most harvested prawn species in Peru, with more than 60 years of commercial fishing history (Reyes-Avalos et al., 2024). As a result, it is also the species most threatened by overexploitation (Wasiw and Yépez, 2017; Pinazo et al., 2021). Other external factors that threaten river prawn populations include habitat destruction, anthropogenic contamination, and destructive fishing practices (Reyes-Avalos et al., 2024). In this context, aquaculture provides an alternative source of seafood that can reduce pressure on wild stocks (Halpern et al., 2021). In Peru, the cultivation of the endemic *M. caementarius* is still in its early stages and is primarily conducted at the research-level (Reyes-Avalos et al., 2024). Additionally, some studies have highlighted the potential for aquaculture of other prawn species including *M. americanum*, mainly due to its large size (Yamasaki-Granados et al., 2013; Pérez-Rodríguez et al., 2018). In this light, our detection assay can be used to verify the species identity of early life stages and juveniles of these prawn species used for restocking and aquaculture purposes through non-invasive sampling methods.

## Conclusions, recommendations, and future research

We presented the development and validation of a proof-of-concept study aimed at the rapid and simultaneous detection of seven commercially valuable Peruvian prawn species using multiplex PCR-SSP technology. Our molecular assay was successfully applied to monitor the trade of cooked samples in restaurants along the Peruvian coast. This study is notable for being the first to provide molecular evidence of the illegal trade of prawns in northern Peru during the closed season. This is particularly significant because current monitoring efforts are primarily directed at the more productive central and southern regions of the country, leaving the northern fishery relatively unmonitored. Further efforts are necessary to establish periodic monitoring programs throughout cities in northern Peru to identify high-priority stages in the supply chain and regions where illegal prawn trade occurs. In this context, our novel multiplex PCR-SSP provides a robust and efficient tool to support monitoring programs, offering accurate, reliable, rapid, and cost-effective identification of Peruvian prawns.

An effective conservation strategy for commercial crustaceans is to impose a permanent ban on ovigerous female prawns. Additionally, enforcing catch limit quotas and designating protected areas based on species-specific assessments would help improve stock health by preventing overfishing and allowing for the reproduction and recovery of the prawn populations. To enhance management and conservation efforts, it would be beneficial to assign unique market names to each prawn species and incorporate molecular identification methods into existing frameworks (Marín, 2025). These measures will improve fisheries and market statistics, facilitate traceability, and help combating mislabeling. Moreover, increased efforts should be directed towards aquaculture production of *M. caementarius* and promoting the trade and consumption of *M. rosenbergii*, particularly during the reproductive closed season of the native species.

Apart from periodic research conducted on *M. caementarius* by Peruvian authorities, there is significant lack of monitoring and population assessments for the other native species. Such studies are urgently needed for these *Macrobrachium* species, as their populations are not only affected by overfishing but also by the predatory threat of feral alien species. For example, nearly two decades ago, a study on the biodiversity of the Tumbes region reported the presence of the exotic Australian redclaw crayfish *Cherax quadricarinatus* (Luque, 2007). Confirmation of the successful establishment and spread of this alien crayfish species was obtained from recent observations of dense populations and collection of multiple specimens in 2024 in freshwater bodies from the Tumbes and Piura regions (R. Alfaro, personal communication, July 23, 2024). Despite that, to date no study has been conducted to assess the impact of these alien crayfish populations on endemic prawns.

Despite existing regulations concerning native Peruvian prawns, the decline in *M. caementarius* populations suggest management deficiencies that require urgent attention. While the wild larvae restocking program of Peruvian *M. caementarius* populations aims to help their populations, the lack of population genetic studies is a concern because these practices can unintentionally reduce genetic diversity, leading to inbreeding depression, decreased fitness, and compromised long-term sustainability of the prawn populations (Grant et al., 2017). In this regard, an ongoing project by the Laboratory of Genetics, Physiology, and Reproduction of the National University of Santa, aims to sequence the complete genome of *M. caementarius*. A high-quality reference genome sequence of this valuable species will enable the development of high-density genomic markers, such as SNPs, which are necessary to assess fine-scale population structure and gene flow (Zhang et al., 2025). Ultimately, this genomic information will significantly aid in future selective breeding programs and germplasm initiatives for this species (Zheng et al., 2024; Zhang et al., 2025).

## Supplementary Figure legends

**Fig. S1**. Simplex and duplex *in-vitro* validation assays. Intraspecific validation PCR results are depicted in Panel 1A for *Macrobrachium caementarius*, Panel 2A for *M. rosenbergii*, Panel 3A for *M. americanum*, Panel 4A for *M. inca*, Panel 5A for *M. gallus*, Panel 6A for *M. panamense*, and Panel 7A for *M. digueti*. Intraspecific validation PCR results are depicted in Panel 1B for *Macrobrachium caementarius*, Panel 2B for *M. rosenbergii*, Panel 3B for *M. americanum*, Panel 4B for *M. inca*, Panel 5B for *M. gallus*, Panel 6B for *M. panamense*, and Panel 7B for *M. digueti*. The labels in the wells for each agarose gel correspond to Ma for *M. americanum*, Mc *M. caementarius*, Md for *M. digueti*, Mg for *M. gallus*, Mi for *M. inca*, Mp for *M. panamense*, and Mr for *M. rosenbergii*. M: molecular ladder (50 bp) loaded into wells at both ends of the agarose gels. NC: negative control.

